# Chimpanzees use advanced spatial cognition to plan least-cost routes

**DOI:** 10.1101/793562

**Authors:** Samantha J. Green, Bryan J. Boruff, Cyril C. Grueter

## Abstract

While the ability of naturally ranging animals to recall the location of food resources and use straight-line routes between them has been demonstrated in several studies, it is not known whether animals can use knowledge of their physical landscape to plan least-cost routes. This ability is likely to be particularly important for animals living in highly variable energy landscapes, where movement costs are exacerbated. Here, we used least-cost modelling to investigate whether chimpanzees (*Pan troglodytes*) living in a rugged, montane environment use advanced cognitive skills to plan energy efficient routes. We used a subset of chimpanzee movement segments together with the available laboratory measurements of chimpanzee energy expenditure to assign movement ‘costs’ which were incorporated in an anisotropic least-cost model and straight-line null model. The least-cost model performed better than the straight-line model across all parameters, and linear mixed modelling showed a strong relationship between the cost of observed chimpanzee travel and predicted least-cost routes. To our knowledge, our study provides the first example of spatial memory for landscape and the ability to plan least-cost routes in non-human animals. These cognitive abilities may be a key trait that have enabled chimpanzees to maintain their energy balance in a low-resource environment. Our findings provide a further example of how the advanced cognitive complexity of hominids have facilitated their adaptation to a variety of environmental conditions and lead us to hypothesise that landscape complexity may play a role in shaping cognition.

## 1 Introduction

Advanced cognitive abilities can facilitate increased foraging efficiency, particularly in animals that depend on clumped resources (Milton, 1981, Benhamou, 1994, Janmaat and Chancellor, 2010, Fagan et al., 2013, Janson, 2019). Research on spatial cognition of naturally ranging animals has generally focused on the abilities of animals to remember the location, type and seasonality of food resources and plan distance-minimising routes between them (Janson and Byrne, 2007, Zuberbühler and Janmaat, 2010). However, it is not known whether animals can remember the physical landscape of their natural environment and use this knowledge to plan energy-minimising routes (Howard et al., 2015).

Physical features of the landscape such as steep slopes or dense vegetation, can significantly increase energy expenditure during foraging (Halsey, 2016) and recent studies have shown that animals will alter their ranging patterns in response to landscape features (Dickson et al., 2005, Wall et al., 2006, Sapir et al., 2011, Newmark and Rickart, 2012, Howard et al., 2015). This landscape driven variation in movement costs is termed the ‘energy landscape’ (Wilson et al., 2012, Shepard et al., 2013) and it follows that animals living in more variable energy landscapes would gain fitness benefits from remembering the landscape of their home range and planning efficient routes. Advances in hand-held Global Positioning Systems (GPS) allowing more accurate collection of movement data in rugged environments and increasingly sophisticated modelling tools that can incorporate landscape features into measures of movement efficiency have opened opportunities for analysis of animal memory of landscape and route choice (Fagan et al., 2013, Kays et al., 2015).

Spatial memory of landscapes and the ability to plan least-cost routes is expected to be most beneficial to animals that a) live in **highly variable energy landscapes**, as the potential savings in movement costs are greater, and b) rely on resources that are (based on Milton, 1981) (i) **stationary**, and therefore predictable in space as opposed to mobile prey, (ii) **patchily distributed**, making random search a less efficient strategy, and (iii) **lower in density**, resulting in increased travel distances between patches and thus increased movement costs.

Chimpanzees (*Pan troglodytes*) have been the subject of more spatial cognition studies than any other non-human primate (Menzel, 2012), but as most field studies have been undertaken in relatively homogenous environments (Wittig and Crockford, 2018), their ability to remember landscapes and incorporate this into their route planning has not been tested. Chimpanzees rely on food resources that are characterised by high spatio-temporal complexity (Janmaat et al., 2016) and thus travel relatively long daily distances, expending more energy on terrestrial locomotion than any other activity (Leonard and Robertson, 1997, Pontzer and Wrangham, 2004). A recent study in Nyungwe National Park, Rwanda showed that chimpanzee ranging patterns can be influenced by their terrain (Green et al., in review), making them perfect subjects to investigate spatial cognition of landscape.

Nyungwe is a low-productivity montane forest (Gross-Camp et al., 2009) in south-west Rwanda that supports a community of chimpanzees that range from 1,795 to 2,951 m ASL, the highest recorded altitude for wild chimpanzees (Green et al., in review). Across locations where chimpanzees have been studied, Nyungwe has one of the most variable energy landscapes, consisting of rugged terrain, dense ground cover and a network of human-made trails that chimpanzees preferentially use for travel (Figure 1, Green et al., in review). The aim of this study was to investigate whether chimpanzees are able to use advanced cognitive skills to reduce energy expenditure in a low productivity environment. We hypothesize that chimpanzees in Nyungwe have spatial memory of landscape and take least-cost routes to goals outside the line of sight (out of sight).

**Figure 1.**
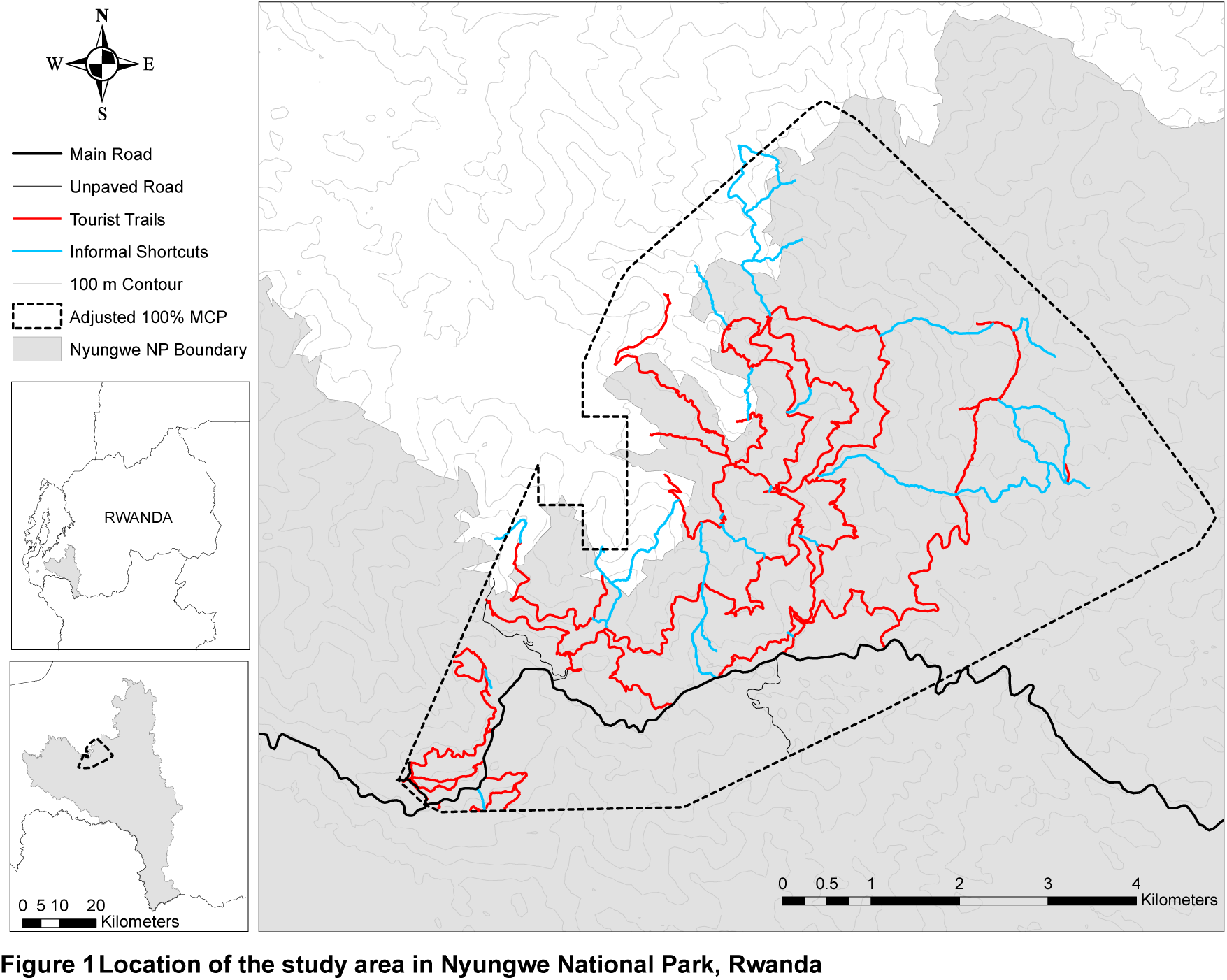
Location of the study area in Nyungwe National Park, Rwanda

These hypotheses were tested using chimpanzee ranging data collected over 14 months and an anisotropic least-cost path (LCP) model that determines the most efficient route (or path of least resistance) assuming full knowledge of the environment. Efficiency is calculated as the cost of moving across the landscape and can include features that impede animal travel, such as slope, vegetation cover or human disturbance (Zeller et al., 2012). Least-cost analysis is increasingly employed to model animal movement for landscape connectivity studies; however, recent reviews found that few studies use empirical data to assign landscape costs or assess model accuracy (Sawyer et al., 2011, Zeller et al., 2012). Most studies also employ isotropic models which are often not realistic (Etherington, 2016), particularly in rugged environments where the cost to travel upslope is greater than downslope for many species (Halsey and White, 2017).

Here, we use change points to define movement phases (Byrne et al., 2009) and use a subset of these movement phases, together with the available laboratory measurements of chimpanzee energy expenditure, to define movement ‘costs’. We then compare the costs and geometry of observed movements with predicted least-cost routes and a straight-line null model to test for evidence of use of cognitive mechanisms to plan energetically efficient routes.

## 2 Methods

Data were collected between November 2016 and December 2017 in Nyungwe National Park, Rwanda (Figure 1). Nyungwe is a rugged montane tropical forest characterised by relatively steep slopes, an open canopy and dense ground vegetation. It protects 1,020 km^2^ of forest and is estimated to contain 380 chimpanzees (IUCN, 2010). The study community range, to our knowledge, to the highest altitudinal limit of their species distribution (2,951 m ASL, Green et al., in review) and consisted of 67 members by the end of the study: 14 adult and 4 sub-adult males, 18 adult and 7 sub-adult females, 12 juveniles and 12 infants (Smith and Green, 2018).

Male chimpanzees are known to travel longer daily distances and have larger home ranges than anestrous females (Wrangham and Smuts, 1980, Chapman and Wrangham, 1993, Doran, 1997, Williams et al., 2002, Bates and Byrne, 2009). Thus, only male chimpanzees were sampled to maximise path data (n = 14 individuals). Focal follows (Altmann, 1974) were undertaken for as long as possible, ideally from nest to nest, on approximately ten days per month. Their locations were recorded at 5 m intervals with a hand-held Garmin GPSMAP 64 device, with GLONASS receiver. The GPS accuracy was within 3–6 m throughout most of the Mayebe home range, but could increase to 20 m in some valleys. Party size and composition was recorded every 15 minutes, with any individuals within 50 m of each other considered to be a part of the same party (following Clark and Wrangham, 1994).

### 2.1 Least-cost analysis

To investigate whether chimpanzees use LCPs when travelling across their environment, we employed the ArcGIS Path Distance Tool which incorporates anisotropy by modifying the cost distance function with a user defined vertical factor. Path Distance calculates the cost of travel between two perpendicularly adjacent cells (*a* and *b*) using the following formula:

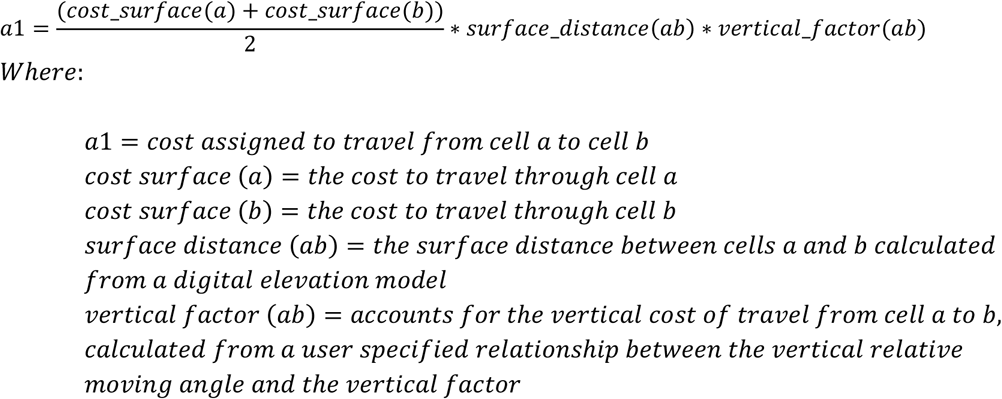

For diagonally connected cells, the larger distance (√2) between cells *a* and *b* is accounted for as follows:

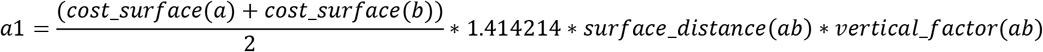

### 2.2 Model inputs

Shepard et al. (2013) identify three landscape factors that influence costs of transport for terrestrial animals: topographic variation, and super and substrate penetrability. Substrate is defined as “the medium over or on which an animal moves” (Shepard et al., 2013 p. 299), and superstrate as “any material against which an animal must push to move” (Shepard et al., 2013 p. 300). Landscape features that had the potential to influence chimpanzee travel in our study area were: slope, trails (reduced superstrate, compact substrate and gently sloping topography), ridges (reduced superstrate) and streams (costly substrate).

A 30 m x 30 m resolution Shuttle Radar Topography Mission (SRTM, available from the US Geological Survey’s EROS Data Center) digital elevation model (DEM) covering the study area was used to calculate a GIS slope layer. Human-made trails were mapped by walking all trails within the Mayebe chimpanzees home range, taking GPS readings every 5 m. This included both tourist trails and informal shortcuts. Established chimpanzee trails were mapped whenever chimpanzees were observed travelling along them. These are trails that were narrower than 1 m, had level substrate that cut into steep slopes, were free from superstrate up to approximately 1 metre (sometimes forming a tunnel through vine thickets), and where bark had been worn off any dead logs or living vines that lay on the trail, suggesting regular use by chimpanzees. These human and established trails were imported into ArcGIS and converted to a raster corresponding to the 30 m x 30 m SRTM DEM.

The ArcGIS 10.6 Hydrology Toolset was used to extract stream and ridge lines from 30 m x 30 m SRTM DEM. Extracted stream and ridge lines were visually inspected using Google Earth and any lines that had not been extracted using automated techniques were manually digitized on screen as described by Gregory et al. (2014). Both stream and ridge lines were converted to a raster corresponding to the 30 m x 30 m SRTM DEM.

### 2.3 Path segmentation

To create daily travel paths, any location points that were less than 30 m apart were discarded to align with the DEM resolution and each consecutive waypoint was then joined with a straight-line segment in ArcMap 10.6 (n = 106 days). The paths were then divided into segments for analyses based on changes in travel direction following Noser and Byrne (2013), Ban et al. (2016), Polansky et al. (2015) and Presotto et al. (2018). We considered a spatial criterion to be more appropriate than a temporal criterion [used by Valero and Byrne (2007) and Bates and Byrne (2009)] as a defined ‘stop time’ would not capture some important determinants of chimpanzee travel routes such as changes in direction after hearing a pant-hoot (fusion) or reaching a tree that bore no ripe fruits (fruit monitoring).

A change point test (CPT) developed by Byrne et al. (2009) was then used to detect significant changes in travel direction. Variants of the test were run from *q* = 1 through *q* = 10 for 10% of the daily travel paths using an alpha level of *p* < 0.05 with *q* = 5 chosen as the most representative since this value maximized the number of change points detected for each day’s path while also failing to ‘overshoot the change point’ (Byrne et al., 2009). After running the CPT on all paths the behaviour associated with each change point was recorded. Change points that were associated with any behaviour other than ‘traveling’ and did not occur on a human-made or established trail, were used to divide paths into segments. Since the CPT would sometimes identify a change point one to two steps away from what could be considered the intuitive change point (Byrne et al., 2009), the behaviour associated with the two location points immediately before and after the detected change point were checked before dividing segments. Any segments less than 150 m were excluded from analysis as movements to out of sight resources was the focus of the study. This resulted in a total of 217 segments.

### 2.4 Movement costs

Fifty segments were set aside for model testing. Whilst it is not possible to isolate the influence of each landscape factor on travel in observational studies, segments chosen for model testing contained travel both off and on trails and incorporated a range of landscapes characteristic from flat to rugged terrain. As there is a paucity of research examining the role of landscape characteristics on energy expenditure of primates, landscape factors were examined in turn to develop a cost surface incorporating topography as well as trails and sub/superstrate.

#### 2.4.1 Vertical factor

To date, the best available information on chimpanzee energetics is Taylor et al.’s (1972) measurements of the energy use of a chimpanzee running on a treadmill with a +15 degree, and −15 degree incline. The chimpanzee used up to 1.75 times more energy on a +15 degree incline and as little as 0.64 times less energy on a −15 degree incline compared to a level surface. As our study area has slopes up to 58 degrees we extrapolated our data based on the trends shown for other quadrupeds (Halsey and White, 2017). As the true function of energy expenditure to slope is not known, we assume a linear function extrapolated for slope ranging from 0 to +58 degrees with values held constant for negative slopes. These values were converted to a Vertical Factor for input to the Path Distance tool (Table 1).

**Table 1.** Extrapolated chimpanzee Vertical Factor Table

| Slope (degrees) | -60 | -45 | -30 | -15 | 0 | 15 | 30 | 45 | 60 |
| --- | --- | --- | --- | --- | --- | --- | --- | --- | --- |
| Vertical Factor | 0.64 | 0.64 | 0.64 | 0.64 | 1 | 1.75 | 2.5 | 3.25 | 4 |

As measured movement costs for chimpanzees are only available for three gradients, a model developed for humans was also tested as recommended by Lempidakis et al. (2018). Tobler’s empirically derived Hiking Function (Tobler, 1993) was used to convert slope to velocity using the following equation:

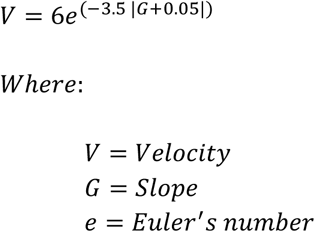

Vertical Factors calculated using this formula (Tripceich, 2009) were used to calculate Path Distance. Both models were tested against segments that did not contain any trail travel. The extrapolated chimpanzee model visually approximated the actual path well, while the human model overestimated the sinuosity of most segments.

#### 2.4.2 Cost surface

##### Trails

Of the remaining landscape features, human-made trails were expected to have the greatest influence on chimpanzee travel patterns. Cost values were iteratively tested for on versus off-trail travel on all test segments that included some trail travel. Relative to the fixed cost of 1, we tested off-trail cell costs from 2 to 10 using 1 increment intervals. As all travel segments were influenced by topography, the segments were tested with and without the Vertical Factors identified previously (Table 1). An off-trail cost value of 2 with Vertical Factors included was the most accurate in predicting the locations where chimpanzees would enter and exit human-made trails.

As the model did not provide a good visual fit for all segments, the cost surface was further refined by including established trails. All cells that contained a human-made trail or established trail were given a value of 1 and all others a value of 2. The majority of segments showed improved visual representation with the addition of established trail costs.

##### Ridges and streams

Adding cost values for ridges and streams did not improve the model for any of the test segments and were therefore excluded from further analysis.

#### 2.5 Final model inputs

The inputs to the Path Distance tool included: a cost surface raster consisting of a 30 x 30 m grid with all human-made and established trail cells representing a cost of 1 and all others a cost of 2; the 30 x 30 m SRTM DEM Surface Raster and the extrapolated chimpanzee Vertical Factor Table (Table 1).

### 2.6 Model accuracy

A Path Distance model was then run for each test segment. The output cost distance and backlink rasters were then used in the ArcGIS Cost Path tool and the cumulative cost for each segment was extracted.

To calculate the cumulative cost of travel on the actual segments, each polyline segment was converted to a 30 m x 30 m raster, and the SRTM DEM cells that corresponded to these rasters were extracted. This was input as a Surface Raster in the Path Distance tool and the same process was followed.

To assess the accuracy of the model, the normalised root mean square error (NRMSE) between the cumulative cost of the actual and modelled segments was calculated by dividing the root mean square error by the range of actual path costs (max_obs_ – min_obs_). The LCP achieved a NRMSE of within 3% (n = 50).

### 2.7 Comparing actual travel to the least-cost model

To test whether chimpanzees use LCPs when travelling in their environment, the costs of actual travel were compared with straight-line travel (the null model) for all remaining segments (n = 167).

To calculate the cumulative cost of linear travel, straight polylines were created between the start and end of each segment in ArcGIS. The polylines were then converted to 30 m x 30 m rasters, and the SRTM DEM cells that corresponded to these rasters were extracted. This was input as a Surface Raster in the Path Distance tool and the same process was followed. The per m costs of actual and modelled paths were calculated by the dividing the cumulative costs by segment length.

To compare the geometry of actual and modelled paths, the sinuosity of each segment was calculated by dividing the least-cost and actual distance by the straight-line distance.

#### 2.7.1 Analysis

The NRMSE was calculated following Howard et al. (2015) to measure how accurately each model predicted actual travel costs and sinuosity. The strength of the relationship between actual travel costs and the least-cost and straight-line models was examined using a linear mixed effects model (LMM) with a Gaussian error structure and identity link function. Actual cost was modelled as the dependent variable and fixed effects were the least-cost and straight-line costs. To account for certain individuals having a disproportionate effect on the dependent variable, the identity of the focal chimpanzee was included as a random effect. To examine potential collinearity among the two independent variables, we determined variance inflation factors (VIF) applied to a standard linear model without the random effects. Multicollinearity was detected in the cumulative cost model (VIF > 10) and cumulative costs were therefore excluded from further analysis. However, the cost per m linear model yielded a VIF of 2.99, which is below recommended cut-offs (Quinn and Keough, 2002, Zuur et al., 2009).

The assumptions of normally distributed and homogeneous residuals were checked by visually inspecting the distribution of the residuals and plotting the residuals against fitted values (Quinn and Keough, 2002). To achieve comparable estimates and increase the likelihood of model convergence, all covariates were z-transformed to a mean of zero and a standard deviation of one before fitting the model (Schielzeth, 2010). To establish the significance of the combined set of predictor variables, we ran a likelihood ratio test comparing the full model with a respective null model containing only the intercept and random effect (Dobson and Barnett, 2002, Forstmeier and Schielzeth, 2011). Marginal (variance explained by the fixed effects) and conditional (variance explained by the entire model, including both fixed and random effects) coefficients of determination were calculated following Nakagawa and Schielzeth (2013) and Johnson (2014).

To test if results changed when the number of alternative routes increases, the same analyses were run for long segments (> 1 km) only (n = 27).

## 3 Results

The least-cost model performed better than the straight-line model across all measures and for all parameters. The cost per m NRMSE’s cumulative and were lowest for the least-cost model, with the straight-line model overestimating both cumulative and per m travel costs (Table 2). The NRMSE between least-cost and actual cumulative costs was less than 5%, indicating that chimpanzee movement segments are similar in ‘cost’ to the least-cost path.

**Table 2.** Costs of chimpanzee travel segments compared to least-cost and straight-line models. SD: standard deviation; NRMSE: normalised root mean square error; Actual: actual path, LCP: least-cost path; Straight: straight-line path.

|  | Model | Cumulative cost |  |  | Cost per m |  |  |
| --- | --- | --- | --- | --- | --- | --- | --- |
|  |  | Mean | SD | NRMSE | Mean | SD | NRMSE |
| All segments<br>(n = 167) | Actual | 954 | 705 | NA | 1.80 | 0.75 | NA |
|  | LCP | 902 | 676 | 3% | 1.72 | 0.71 | 6% |
|  | Straight | 1,150 | 1,024 | 10% | 2.23 | 0.83 | 17% |
| Segments > 1 km<br>(n = 27) | Actual | 2,079 | 884 | NA | 1.37 | 0.33 | NA |
|  | LCP | 1,967 | 869 | 4% | 1.35 | 0.31 | 8% |
|  | Straight | 2,757 | 1,386 | 28% | 2.23 | 0.58 | 57% |

The full model with the least-cost and straight-line costs as predictor variables and actual costs as the response, was significantly different from the null model containing only the intercept and the random effect (chi sq = 100.84, df = 3, p = <0.001). The interaction between least-cost and straight-line models was not significant and the model was thus rerun without the interaction effect. The final model revealed a significant effect of both the least-cost and straight-line models on actual cost (Table 3), but separate models revealed that the least-cost model explained 91% of the variation in actual costs (estimate = 0.71, standard error = 0.02, r^2^ 0.91, p = <0.001, Figure 2), while the straight-line model only explained 66% of the variation (estimate = 0.60, standard error = 0.3, r^2^ 0.66, p = <0.001, Figure 2).

**Table 3.** Results of the LMM with actual costs per m as the response variable (n = 167)

| Predictor Variable | Estimate | Std. Error | t | 95% CI | P |
| --- | --- | --- | --- | --- | --- |
| Intercept | 1.79 | 0.02 |  |  |  |
| LCP | 0.65 | 0.03 | 22.07 | 0.02, 0.14 | <0.001 |
| Straight | 0.08 | 0.03 | 2.67 | 0.59, 0.71 | 0.008 |

**Figure 2.**
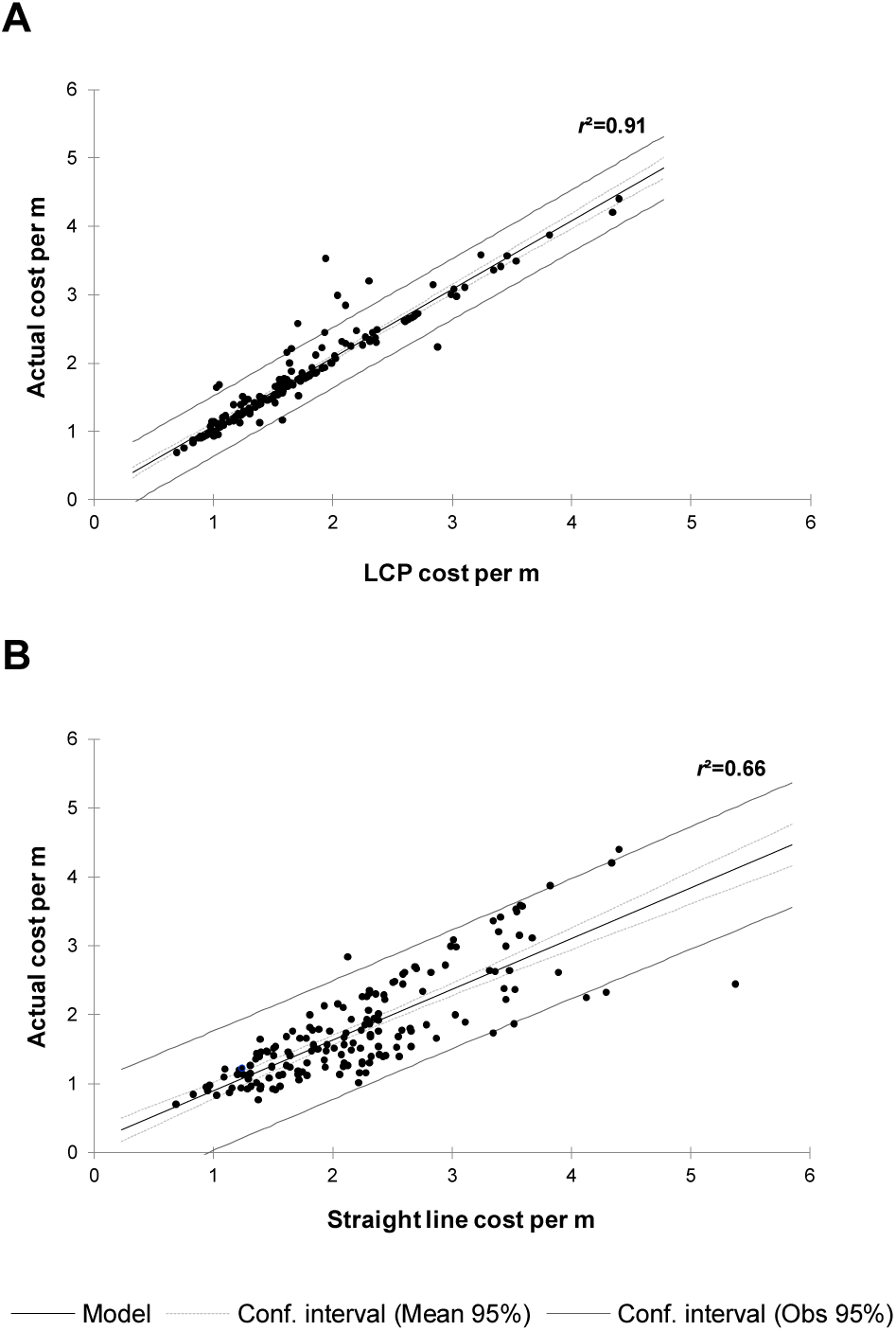
Relationship between the actual cost per m and the least-cost model (A),and straight-line model (B) for 167 travel segments

These results held true for distant goals (>1 km in length). The NRMSE between actual and least-cost cumulative costs remained less than 5% and the NRMSE between per m costs remained less than 10%, while the NRSME between actual and straight-line costs more than doubled for both cumulative and per m costs (Table 2). The LMM showed that the least-cost model was still a strong predictor of actual per m costs for long segments (estimate = 0.31, standard error = 0.02, *r*^2^ = 0.88, p = <0.001).

The least-cost model is also a better predictor of actual travel sinuosity and length than the straight-line model (Table 4), with chimpanzees taking longer, more sinuous paths that incorporate trails and/or avoid steep inclines (e.g. Figure 3).

**Table 4.** Length and sinuosity of chimpanzee travel segments compared to least-cost and straight-line models. SD: standard deviation; NRMSE: normalised root mean square error; Actual: actual path, LCP: least-cost path; Straight: straight-line path.

| Model | Length (m) |  |  | Sinuosity |  |  |
| --- | --- | --- | --- | --- | --- | --- |
|  | Mean | SD | NRMSE | Mean | SD | NRMSE |
| Actual | 596 | 519 | NA | 1.12 | 0.14 | NA |
| LCP | 592 | 509 | 2% | 1.09 | 0.12 | 13% |
| Straight Line | 523 | 431 | 4% | 1 | 0 | 22% |

**Figure 3.**
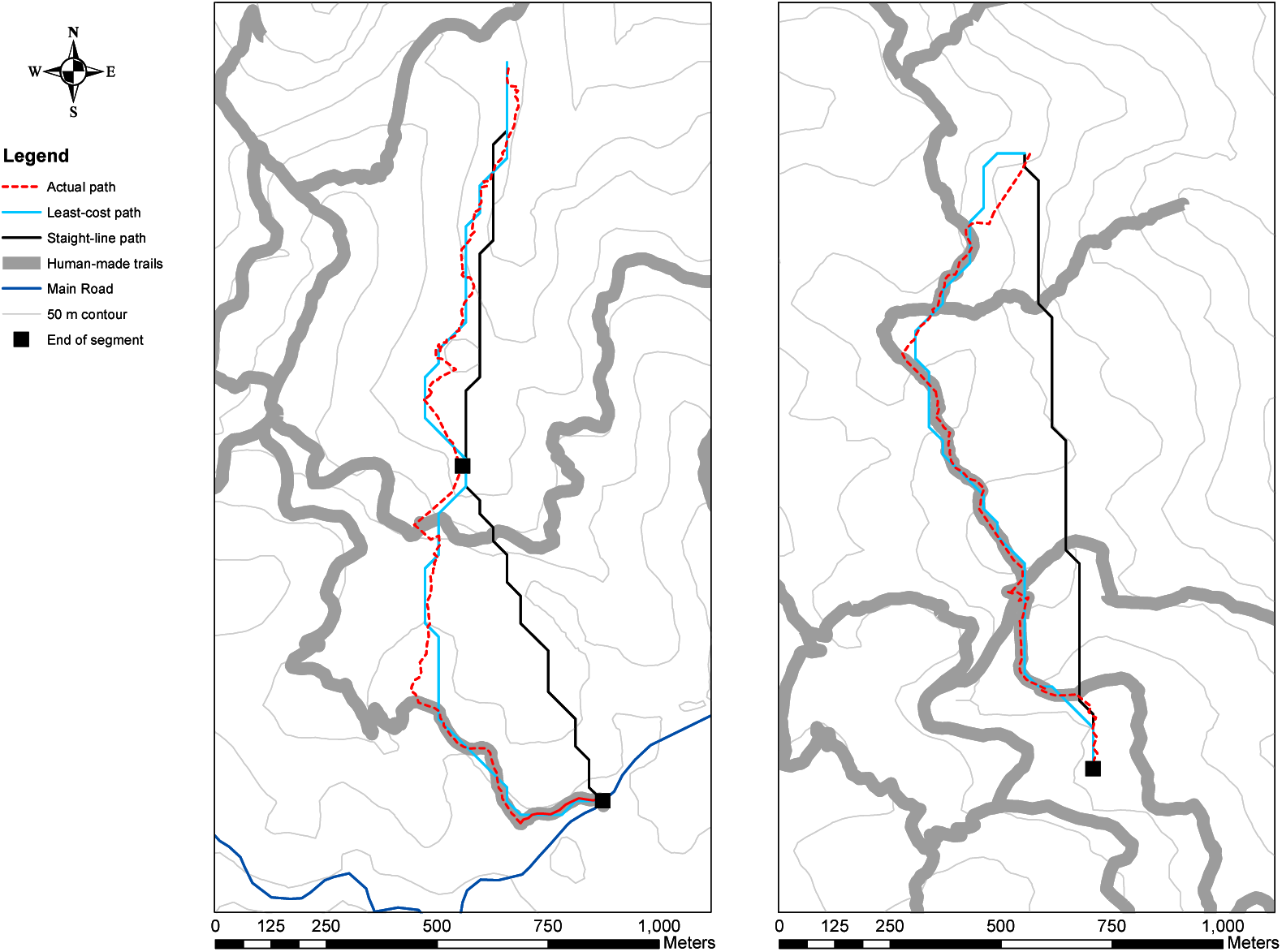
Modelled and actual path segments on 9 April 2017 (left) and 27 November 2017 (right)

## 4 Discussion

The ability of naturally ranging animals to remember the location of food resources and plan least-distance routes has been demonstrated in several studies (review in Trapanese et al., 2018), but the extent to which animals remember the landscape of their home range and plan least-cost routes is not well understood. By using chimpanzee ranging data and anisotropic least-cost modelling we were able to investigate chimpanzee spatial memory of landscape and test their ability to plan efficient foraging routes in a variable energy landscape.

The least-cost model predicted the costs and sinuosity of chimpanzee paths better than the straight-line model and linear mixed modelling showed a strong relationship between the costs of chimpanzee travel and the modelled least-cost routes. These results cannot be explained by use of visual cues, as the minimum segment length (150 m) was substantially greater than the distance from which human observers could see trails and other prominent landmarks in Nyungwe (S.J. Green, unpublished data) and aligns with the visual perception radius used in tests of chimpanzee spatial memory of fruit trees in a previous study (Janmaat et al., 2013). Whilst the travel costs do not represent metabolic rates, the available laboratory data on chimpanzee energetics was used to inform vertical factor calculations and the cost surface was calibrated to a subset of chimpanzee pathway data, which recent reviews have recommended as the most ecological meaningful technique (Zeller et al., 2012, Etherington, 2016). Outputs not based on modelled costs (travel segment sinuosity and length) also showed better agreement with the least-cost than the straight-line model. As the model employed assumes complete knowledge of the landscape (Etherington, 2016), our results provide strong evidence that Nyungwe chimpanzees have comprehensive spatial memory of their home range landscape and plan least-cost routes to out of sight goals. Chimpanzees demonstrated remarkable spatial accuracy in planning least-cost routes, even for long (>1 km) movement segments when the number of potential alternative routes increases considerably. These results differ to previous studies which found no relationship between predicted least-cost and actual travel routes in non-human primates (Gregory, 2011, Howard et al., 2015). However, these findings are likely due to the use of unrealistic isotropic models (Etherington, 2016) and lack of model calibration to pathway data (Zeller et al., 2012) and not a reflection of the animals’ cognitive abilities.

Demonstrating spatial knowledge in naturally ranging animals that travel in relatively linear segments is difficult, as straight-line movement can be associated with a number of foraging processes that are not goal-orientated (Janson and Byrne, 2007). Thus, a number of onerous measurements are required to infer cognitive processes (Janson and Byrne, 2007), such as recording all alternative food resources bypassed (Normand et al., 2009, Janmaat et al., 2013), and identifying which of those resources are more valuable, which can be extremely difficult in itself (Ban et al., 2016). Our work demonstrates that least-cost path modelling can offer an alternative approach to assess cognitive abilities in wild animals that are known to modulate their movements in response to energy landscapes.

Ecological models, by their nature, represent a simplified version of the natural environment and are therefore limited in their ability to capture the full complexity of interactions between landscape features and animal movement. Whilst our model was able to predict the travel costs of chimpanzees within a 3% error, analysing where the model did not fit well can yield important insights into other key drivers of animal movement (Shepard et al., 2013, Lempidakis et al., 2018). Some inconsistencies could be explained by the lack of a detailed super and substrate map, which resulted in landscape features that facilitate chimpanzee movement (e.g. exposed rocks, vines that allowed chimpanzees to climb up or down steep cliff faces and fallen logs that enable stream crossings) being omitted from the cost surface. The importance of compact substrate and reduced superstrate was demonstrated by one occasion when the focal chimpanzee deviated 75 m from the predicted least-cost route to travel along an area at the altitudinal limit of their home-range with exposed rocks and sparse ground cover. Additionally, the DEM used was coarser in resolution than other model elements (e.g. trails). This sometimes resulted in the least-cost model underestimating actual travel costs. More detailed elevation, super and substrate layers could be obtained using high resolution imagery collected using new satellite constellations (Planet Labs), or surveys flown by manned and Unmanned Aerial Vehicles (UAV)’s. Landscape features could be extracted with the aid of LiDAR classification softwares to produce high resolution cost surfaces (see Strandburg-Peshkin et al., 2017 for an example). With fine-scale remote sensing technologies becoming increasingly more affordable, this offers an exciting area for future research.

To our knowledge, this study provides the first example of spatial memory for landscape and the ability to plan least-cost routes in non-human animals. As the energetic cost of terrestrial locomotion comprises a substantial proportion of a chimpanzees daily energy budget, the ability to plan energy-minimising routes may be a key trait that has allowed Nyungwe chimpanzees to survive in a low-resource, montane environment. Our study provides a further example of how the advanced mental complexity of hominids may have facilitated their adaptation to a variety of environmental conditions.

Recent research has renewed interest in the role ecological variation plays in shaping cognitive abilities (Rosati, 2017). The ‘ecological intelligence’ or ‘harsh environment’ hypothesis argues that environments with resources that are low in abundance, sparsely distributed and ephemeral, favour the development of mental abilities that facilitate efficient foraging (Milton, 1981, Milton, 1988, Sol et al., 2005, Dukas, 2009). There is growing evidence to support this from comparative studies of primates (review in Rosati, 2017), birds (Freas et al., 2012, Pravosudov and Roth, 2013, Roth et al., 2013, Sonnenberg et al., 2019), bats and rodents (review in Harvey and Krebs, 1990). However, a recent study highlighted the need for a clearer definition of what constitutes environmental ‘harshness’ (Hermer et al., 2018). Based on the results of this study, we propose that environmental harshness could be expanded beyond the spatio-temporal complexity of food resources to include landscape complexity. Studies in relatively homogenous landscapes have provided good evidence that chimpanzees consider the Euclidean distances between potential food trees when deciding where to forage (Normand et al., 2009). Chimpanzees in highly variable energy landscapes face the additional cognitive load of recalling the landscape between themselves and potential food trees and comparing the least-cost routes between them. Future research on the cognitive abilities used by chimpanzees and other large-brained animals to navigate a variety of landscapes is required to shed light on the role energy landscapes play in shaping animal cognition.

## 5 Conclusion

While there is growing evidence that naturally ranging animals are able to remember the type and location of food resources (Fagan et al., 2013, Trapanese et al., 2018), their ability to plan efficient foraging routes in heterogenous landscapes is not well understood. By using empirical data to define chimpanzee movement costs and anisotropic least-cost modelling, we provide the first evidence for spatial memory of landscape and planning of least-cost routes by a non-human animal. These cognitive abilities may be key to chimpanzee survival in low resource, montane environments and may have been shaped by the ‘harshness’ of their energy landscape. Application of least-cost modelling in cognitive studies of other naturally ranging animals in a variety of landscapes would shed light on this.

## Acknowledgments

We thank the Tony Mudakikwa and Albert Kayitare at RDB for permission to conduct research at Nyungwe National Park, Innocent Ndikubwimana and Kambogo Ildephonse at RDB for facilitating our research, the RDB and Wildlife Conservation Society trackers for their assistance in locating the chimpanzees, and Donat Murwanashyaka and Brad Smith for their assistance in chimpanzee tracking and data collection. The research conducted adhered to the legal requirements of Rwanda and complied with all Rwanda Development Board regulations. Funding was provided by the University of Western Australia and Basler Stiftung für biologische Forschung.

## Author contributions

S.J.G. collected and analysed data, S.J.G, B.J.B and C.C.G wrote the paper.

